# Contact area and tissue growth dynamics shape synthetic juxtacrine signaling patterns

**DOI:** 10.1101/2023.07.12.548752

**Authors:** Jonathan E. Dawson, Abby Bryant, Trevor Jordan, Simran Bhikot, Shawn Macon, Breana Walton, Amber Ajamu-Johnson, Paul D. Langridge, Abdul N. Malmi-Kakkada

## Abstract

Cell-cell communication through direct contact, or juxtacrine signaling, is important in development, disease, and many areas of physiology. Synthetic forms of juxtacrine signaling can be precisely controlled and operate orthogonally to native processes, making them a powerful reductionist tool with which to address fundamental questions in cell-cell communication *in vivo*. Here we investigate how cell-cell contact length and tissue growth dynamics affect juxtacrine signal responses through implementing a custom synthetic gene circuit in *Drosophila* wing imaginal discs alongside mathematical modeling to determine synthetic Notch (synNotch) activation patterns. We find that the area of contact between cells largely determines the extent of syn-Notch activation, leading to the prediction that the shape of the interface between signal-sending and signal-receiving cells will impact the magnitude of the synNotch response. Notably, synNotch outputs form a graded spatial profile that extends several cell diameters from the signal source, providing evidence that the response to juxtacrine signals can persist in cells as they proliferate away from source cells, or that cells remain able to communicate directly over several cell diameters. Our model suggests the former mechanism may be sufficient, since it predicts graded outputs without diffusion or long-range cell-cell communication. Overall, we identify that cell-cell contact area together with output synthesis and decay rates likely govern the pattern of synNotch outputs in both space and time during tissue growth, insights that may have broader implications for juxtacrine signaling in general.

## INTRODUCTION

Juxtacrine or contact-dependent signaling involves a signal-sending cell displaying a membrane-anchored ligand that engages a receptor on the surface of an adjacent signal receiving cell^1,2^. This leads to a signal response, such as a change in cell behavior important for morphogenesis, inflammation, tissue repair, or a host of other biological processes^2–5^. Synthetic cell-cell communication, where biological and engineering principles are combined to control tissue behavior, is emerging as a future biomedical tool that could allow natural developmental programs governing growth, differentiation, or cell death to be deployed at customized times or locations^6–8^. The approach also offers an opportunity to elucidate the principles governing the development of tissues^9^ as insights can be gained through constructing synthetic circuits that mimic native programs to spatially pattern cellular differentiation and growth^5,10–13^.

A key challenge in synthetic biology is to engineer signaling systems to create an output at precise times and locations, such that cells gain positional information and regulate cell-fate decisions. However, such patterning is a complex process that depends on factors such as cell surface to volume ratio, contact curvature and the overall cell volume^14,15^, all of which have the potential to alter a juxtacrine signal in a rapidly growing tissue^14,16,17^. Recent studies have successfully implemented synthetic patterning circuits in bacteria and cells in culture^12,18^. An important next step is to understand how the changes in cell number and shape associated with tissue growth might alter the engineered responses. To address this, we have produced a synthetic juxtacrine system for use in the developing *Drosophila* wing imaginal disc, together with a minimal mathematical model of its operation within a field of proliferating cells. The system is based on synthetic versions of the Notch receptor (called synNotch), which has the capacity to receive a unique juxtacrine signal and produce a customizable output^19,20^. We find that a larger area of contact between signal sending and receiving cells correlates with enhanced synNotch activation, leading us to predict that the shape of the interface between them will markedly alter the extent of the output *in vivo*. We also present evidence that juxtacrine output coupled with cell division sets up an emergent graded spatial output profile. Rather than being restricted only to the synNotch cells adjacent to ligand cells, the output can extend several cells away from the interface *in vivo* and our model indicates that this depends on the rates of output biosynthesis and degradation, together with cell division. These results highlight considerations that will be important in engineering reproducible cell behavior *in vivo* and that native juxtacrine pathways must counter to elicit precise positional information.

## RESULTS

### A synNotch circuit in growing *Drosophila* epithelia leads to heterogeneous output patterns

The *Drosophila* wing imaginal disc is a long-standing and powerful model epithelial tissue within which cell fate patterning and growth are exceptionally well understood^21–23^ and with established techniques for controlling and monitoring signaling events^24,25^. It undergoes extensive proliferation^26^ and shape changes^27^ making it ideal to study the impact of tissue geometry and growth on the output of a juxtacrine signal (referred to here simply as output). During development, the wing disc grows from ∼5-40 cells to ∼40,000 cells within a time span of 3 days, at the end of which signaling events can be conveniently measured at the third instar stage^28^. Synthetic cell-cell communication provides a simplified juxtacrine signaling system for analysis *in vivo* since the signal inputs and outputs can be customized and operate orthogonally to endogenous signal systems that govern normal tissue growth^29,30^.

Given these advantages, we developed a synthetic juxtacrine signaling system based on the Notch receptor, an important and pervasive signaling system in the development of all animals^31,32^. The Notch receptor and its ligands, such as the *Drosophila* ligand Delta, are single-pass transmembrane proteins. Delta, presented on the cell surface, binds to the ectodomain of Notch on neighboring cells and this physical interaction activates the receptor by inducing proteolytic cleavage of Notch within the extracellular Notch Regulatory Region (NRR), which is followed by an intramembrane *γ*-secretase cleavage. This results in the intracellular domain moving to the nucleus to act as a transcriptional activator of Notch target genes, such as *cut* (as described in detail, Fig 1(A)). We produced a functional synNotch receptor by replacing three functional domains of native Notch with heterologous counterparts: 1) The ligand binding epidermal growth factor (EGF) repeats of Notch were replaced with part of the extracellular domain of the follice-stimulating hormone (FSH) receptor. We have previously shown that a receptor with this change (FSHR-Notch) is activated by FSH-Delta, a chimeric Delta where the extracellular domain is replaced with Follicle Stimulating Hormone. 2) The NRR was replaced with the mechanosensitive A2 cleavage domain from the blood clotting protein von Willibrand Factor, previously shown to recapitulate NRR function, albeit with a distinct level of receptor activation suitable for this study^33,34^. 3) In place of the Notch intracellular domain, we introduced a heterologous transcription factor domain used routinely in a *Drosophila* bipartite expression system, LexA-VP16. These three changes produced FSHR-synNotch, in which only the transmembrane domain derives from native Notch. The coding sequences of both FSH-Delta and FSHR-synNotch were placed downstream of an *Upstream Activating Sequence* (*UAS*)-dependent promoter controlled by the yeast Gal4 transcription factor^35^ and paired with the *nubbin*.*Gal4* (*nub*.*Gal4*) transgene to drive the expression of *UAS>FSRH-Notch* and *UAS>FSH-Delta* transgenes in the developing wing imaginal disc (referred to here as wing).

**FIG. 1:**
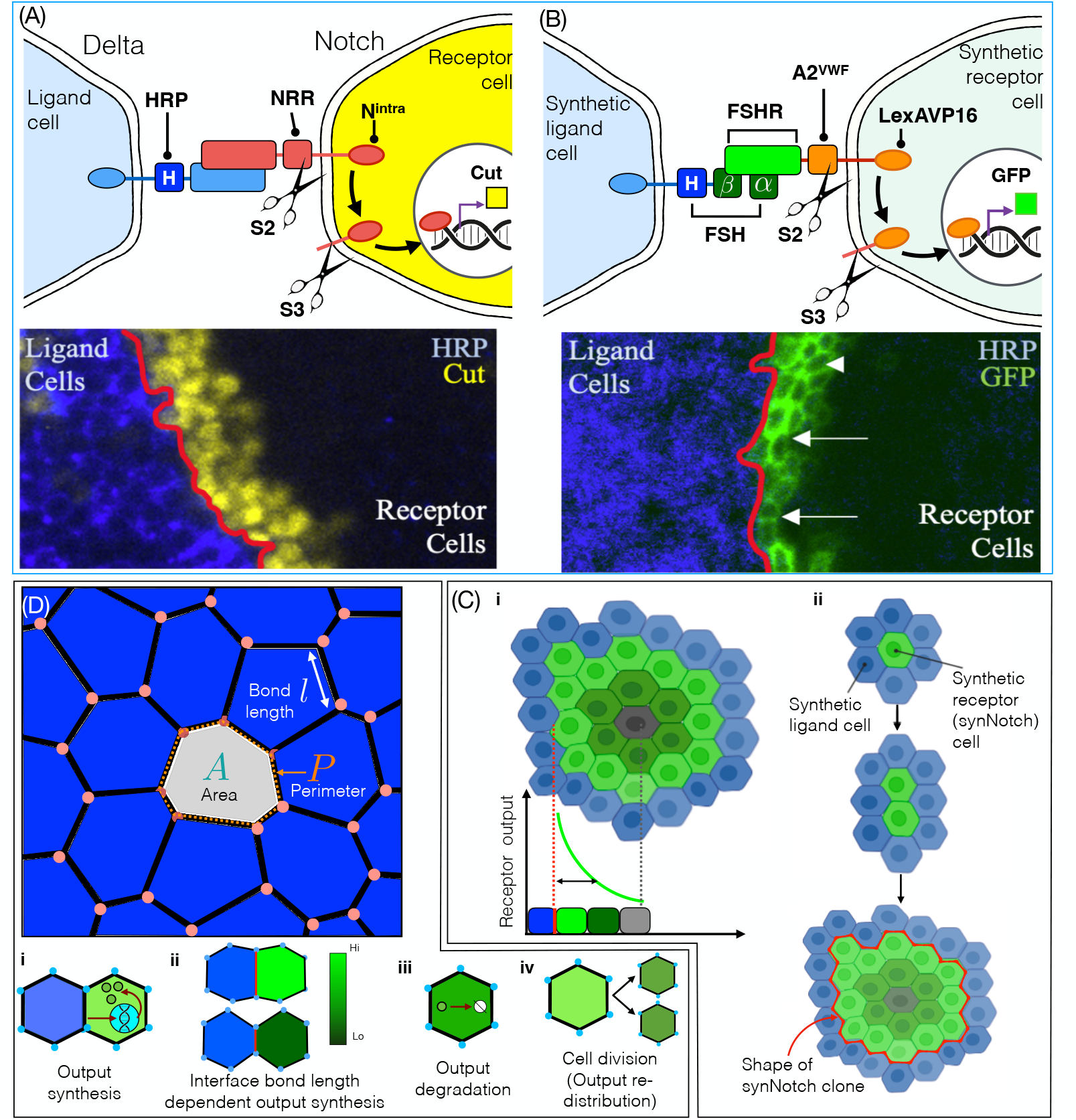
A combined *in vivo* and computational toolkit reveals marked spatial heterogeneity in a synthetic juxtacrine signal. **(A)** Top: Binding of ligand, such as *Drosophila* Delta, to Notch induces S2 cleavage of the Negative Regulatory Region (NRR), S3 cleavage of the transmembrane domain, nuclear access of the intracellular domain (N^intra^), and activation of target genes (e.g. *cut*). Bottom: Confocal image of ectopic Cut (yellow) in the nuclei of *Drosophila* wing disc cells overexpressing Notch adjacent to HRP tagged Delta expressing cells (blue). **(B)** Top: The synNotch activation mechanism is the same as for native Notch (A), although the following components differ: The Delta extracellular domain was replaced with the *β*-subunit of follicle stimulating hormone, and FSH*α* is co-expressed to reconstitute the composite FSH ligand (FSH-Delta); the Notch ligand binding region was replaced by the FSH receptor ectodomain; the NRR was replaced with the A2 domain force sensor from von Willibrand factor; and N^intra^ was replaced with the lexA-VP16 transcription factor. This FSHR-synNotch receptor activated the lexAOP rCD2-GFP transgene (referred to here as GFP). Bottom: Confocal image of GFP output at the surface of FSHR-synNotch cells (uncolored) that are adjacent to HRP tagged FSH-Delta cells (blue). The intensity of GFP varies (arrows) and extends several cell diameters from the clone interface (arrowhead). **(C)** Graphic depiction of how a single synNotch cell surrounded by ligand cells can divide to become a clonal population (clone) of FSHR-synNotch cells with a distinct level and pattern of output, along with a unique clone shape. **(D)** 2D computational vertex model for synNotch induced gene expression. Cells are represented by polygons composed of vertices (orange dots) with a preferred area A_0_ and perimeter P_0_. The cell shape is characterized by an energy function that describes the balance between cell area compressibility and cell perimeter contractility (Materials and Methods Section 2) and the major parameters are (i-ii) cell-cell contact length dependent output synthesis rate (S), (iii) output degradation rate (D) and (iv) the rate of cell division, upon which the GFP output in the parent is redistributed equally between the two daughter cells (iv).

The Mosaic Analysis by Promoter Swap (MAPS) technique^34^ was used to induce mitotic recombination between *UAS>FSRH-Notch* and *UAS>FSH-Delta* transgenes early in development (see Materials and Methods, Section 1), the outcome of which is a clonal population (or clone) of signal sending FSH-Delta cells (referred to here as ‘ligand cells’) and signal receiving cells that express FSHR-synNotch (‘synNotch cells’). Thus signaling interfaces are produced in the wing disc where cells expressing FSH-Delta meet adjacent cells that express FSHR-synNotch, within which the FSHR-synNotch transcriptional response or ‘output’ can be assessed. Expressing these components does not change Cut expression nor the development of the wing (Supplementary Information (SI) Section 1 and Fig. S1), indicating that FSHR-synNotch operates orthogonally to native Notch and other signaling pathways.

To visualize FSHR-synNotch activation, we included a LexA-VP16 responder transgene composed of the LexOp promoter upstream of the coding sequence of a surface localized CD2-GFP chimeric protein (*LexAOP*.*rCD2-GFP* (see Fig 1(B)). In the wing, we observed native fluorescence of this GFP output primarily within synNotch cells adjacent to ligand cells, but also within synNotch cells a few cell diameters from the ligand-synNotch cell interface (arrowhead in Fig. 1(B)). Moreover, the GFP intensity along the interface was not uniform (cells with different intensities are marked with arrows in Fig 1(B)). This suggests that in proliferating epithelial tissues the FSHR-synNotch response is variable and not restricted to those synNotch cells in direct contact with ligand cells as might be expected with a juxtacrine signal. Therefore to better understand these signal output patterns, and juxtacrine communication within proliferating tissues more generally, we developed a novel minimal computational model described below.

### A new computational model predicts that synNotch signal output depends on cell-cell contact length

Tissues in developing organisms are growing active materials with variable geometrical and mechanical properties^36–39^. In particular, cell division-driven changes in the number and spatial arrangement of synNotch and ligand cells can dramatically alter the contact interface between them (Fig. 1(C i,ii)) and impact output patterns. In signal receiving cells co-cultured with signal sending cells, Notch activation increases with the area of contact between the two cell types^40^. To directly test whether synNotch output *in vivo* correlates with the contact area between synNotch and ligand cells, we developed a vertex-based^41,42^ mathematical model with mechanochemical coupling between cell-cell contact length and synNotch output (see Fig. 1(D) and Materials and Methods Sections 2-4 for details).

We predicted that the synNotch output (*G*_*α*_ [arbitrary units] as a function of time (*t*) representing GFP) increases with the contact length between synNotch and ligand cells (Fig. 1(D-ii))^40^ and the rate of GFP synthesis (*S*) (Materials and Methods Section 3). Our model considers two cell types: (i) synNotch cells (initially with no output (Fig. 1(D), gray) which produces GFP upon synNotch activation (Fig. 1(D-i), green)) and (ii) ligand cells that activate synNotch in adjacent cells only (Fig. 1(D), blue). The output strength in synNotch cells is visualized by a green color scale, with dark green denoting low output and bright green color representing high output (Fig. 1(D-ii)). In each cell, GFP is continuously degraded at the rate of *D* [per hour (*h*^−1^)] (Fig. 1(D-iii)). We simulate tissue growth by allowing each cell to grow in size until its area has doubled upon which it divides along its short axis with the GFP output distributed equally between the two daughter cells (Fig. 1(D-iv)) (see Materials and Methods Section 4 for details and Table (1) for relevant simulation parameters).

We first used the model to predict receptor activation patterns following repeated cycles of cell growth and division over a time period similar to that used to produce clonal signaling populations in the wing. The model starting condition is a single synNotch cell surrounded by ligand cells and for several cell divisions all synNotch cells produce a GFP output. Following proliferation equivalent to 50-80 hours, a ‘donut’ pattern emerges where the output intensity is higher in the synNotch cells at the clonal interface compared to the synNotch cells in the clone interior (S1 Movies 1-3, Fig. 2(A)). To verify this prediction *in vivo*, we imaged clones in the wing after 24, 48 and 72*h* of proliferation. At the first two time points all synNotch cells were in contact with ligand cells and produced a GFP output. After 72 hours the synNotch clone was larger and cells in the clonal interior lacked detectable GFP expression (Figs. 2 (B-D)). Both these observations are consistent with the model predictions and indicate that our model can successfully predict patterns of receptor activation in the wing as populations of ligand and synNotch cells grow and divide.

**FIG. 2:**
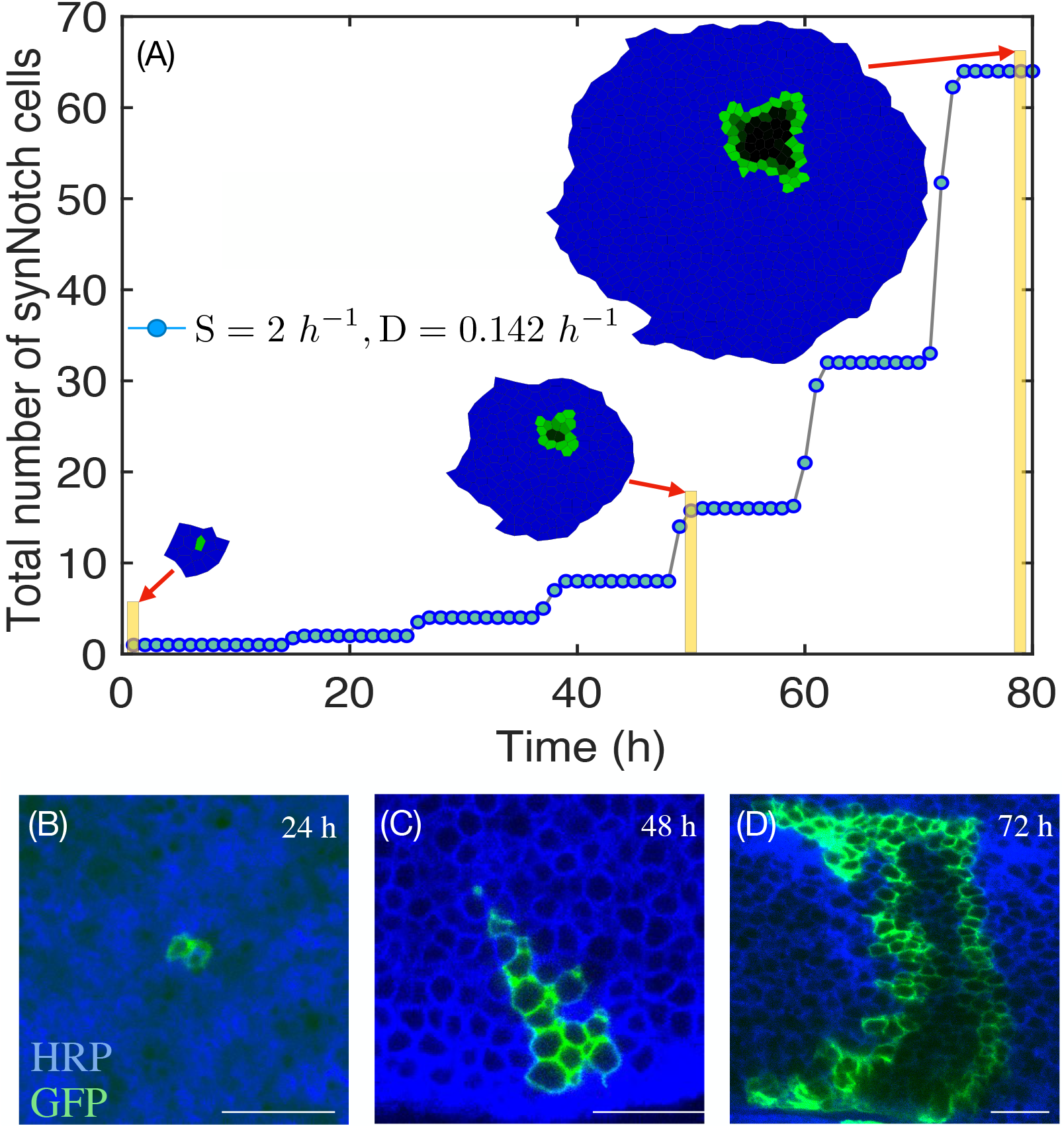
Cell division leads to distinct clone shapes and emergent differences in synNotch output patterns. **(A)** Simulation of a field of proliferating cells with one initial synNotch cell (uncolored) surrounded by ligand cells (blue) at the initial time (*t*_0_). Output synthesis (S) and degradation (D) rates are fixed. As cell number increases over time, the clonal synNotch populations take on distinct clone shapes with unique output patterns (see inset). **(B-D)** Clonal populations within wing discs at 24, 48 and 72 hours after generation of a single FSHR-synNotch cell surrounded by FSH-Delta cells (blue) at *t*_0_. At 24 hours, all synNotch cells express GFP (green), but after 72 hours GFP is not detected in the interior of the synNotch clone, a trend that qualitatively resembles the simulated patterns (A). Each scale bar represents 10*μm*.

Next, we examined the importance of ligand-synNotch cell contact area in producing the synNotch output, approximated in the two-dimensional (2D) model by the contact length between synNotch and ligand cells. Our model predicted that the contact length changes extensively following repeated rounds of growth and cell division, as shown for a typical ligand-synNotch interface (Fig. 3(A)). In this example, some synNotch cells are in contact with a single ligand cell whilst others are in contact with as many as four distinct ligand cells, depending on the synNotch cell’s shape and position within the clonal population. In our equation governing the dynamics of GFP output (See Eq.(3) in Materials and Methods Section 3), the term representing GFP synthesis depends on a base rate of GFP synthesis, *S*, and the contact length between ligand-synNotch cells. Consequently, our model predicts a strong linear dependence between the cell-cell contact length, which can vary from 0 − 15μm, and the output intensity (Fig. 3(B)) with larger contact length leading to higher output signal intensity.

**FIG. 3:**
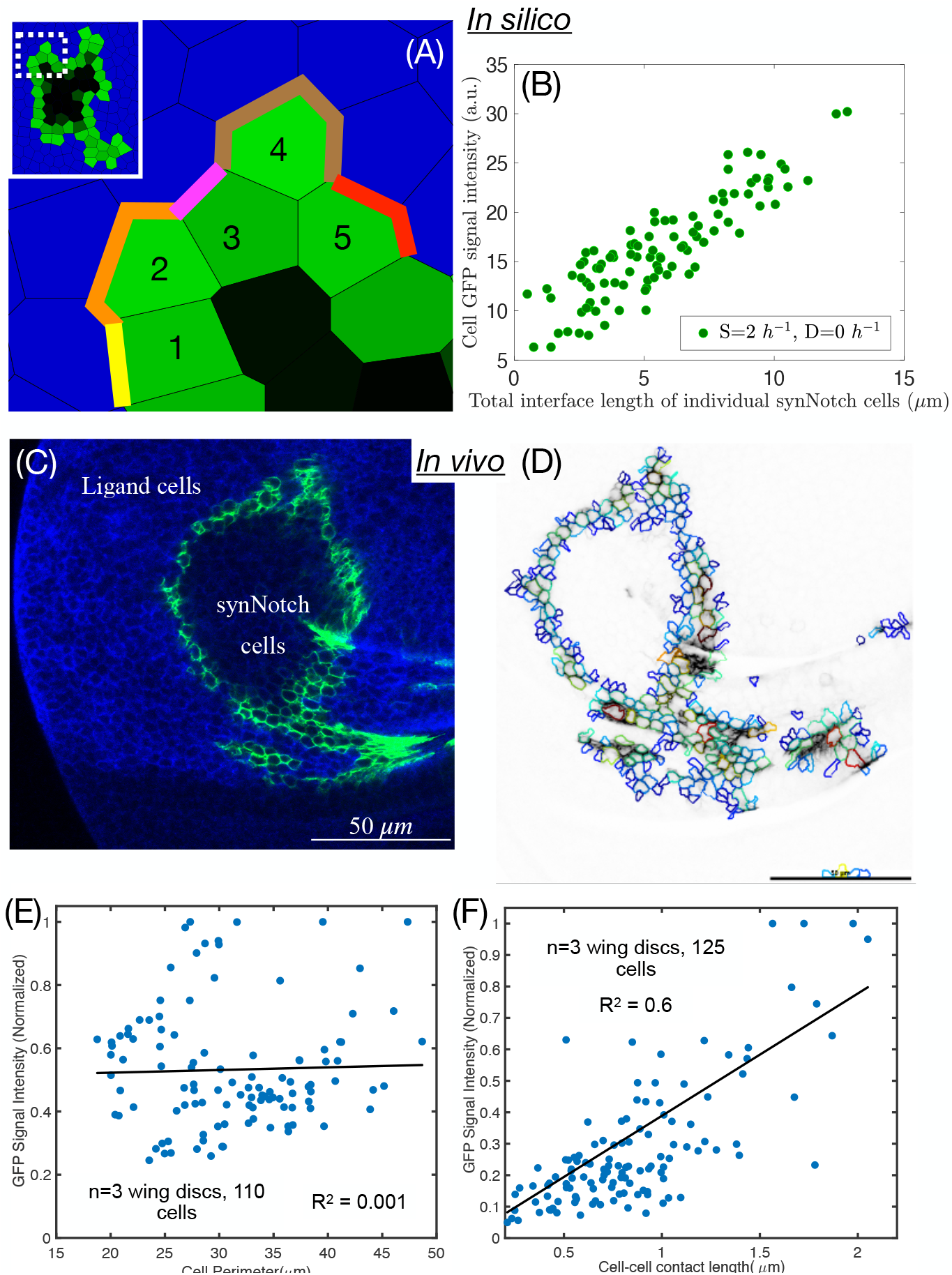
Cell-cell contact length affects synNotch output. **(A)** There is extensive variation in cell-cell contact length which is more pronounced at points of inflection (boxed region, inset) in synNotch clones, e.g. cell #1 has a short contact length with ligand cells while cell #4 has a longer contact length. **(B)** Output intensity in the border synNotch cells increases with the total contact length between synNotch and ligand cells. **(C**,**D)** Cell segmentation analysis (D) was generated using Cell Profiler43,44 from a wing disc (C) with a synNotch clone (expressing GFP) surrounded by ligand cells (blue) to obtain cell perimeter values vs integrated output intensities (see Materials and Methods, Section 5). **(E**,**F)** Output does not depend on the size of individual cells (E, 110 synNotch cells from 3 wing discs) but does increase proportionally with contact length (F, 125 synNotch cells from 3 wing discs).

To test this prediction, we analyzed *in vivo* images of wing disc cells with synNotch response (see Fig. 3(C-D)). We compared the total GFP signal intensity to synNotch cell perimeter and found no correlation (Fig. 3(E)) indicating that cell size *per se* is not predictive of synNotch output. In contrast, GFP output shows a strong correlation with the ligand-synNotch cell-cell contact length with a linear trendline in Fig. 3(F). This strongly supports our prediction that longer contact lengths result in higher synNotch output. Overall, our model focusing on cell-cell contact length dependent synNotch activation can successfully recapitulate time-dependent patterns of receptor activation in a complex, rapidly proliferating tissue.

### Heterogeneous output along the synNotch clonal boundary is due to large variations in cell-cell contact length

In both our model and *in vivo* images, we noted marked variations in the GFP expression in synNotch cells at the signaling interface (as in SI Fig. S2(A-B)). We hypothesized that the cell-cell contact area variations lead to this heterogeneous synNotch output. To test this, we first quantified the GFP output of synNotch cells along the interface *in silico* (see Fig. 4(A)). To do this consistently, we assigned angular positions *θ*_*k*_ to border synNotch cells with the center of the synNotch clone marked with pink dot in Fig. 4(A). In general, we observe large fluctuations in the GFP intensity in cells along the border as observed from the peaks and troughs in Fig. 4(B). The fluctuation amplitude depends on the output decay rate (*D*), although at all values of *D*, the fluctuations persist (Fig. 4(B) and Materials and Methods, Section 3).

**FIG. 4:**
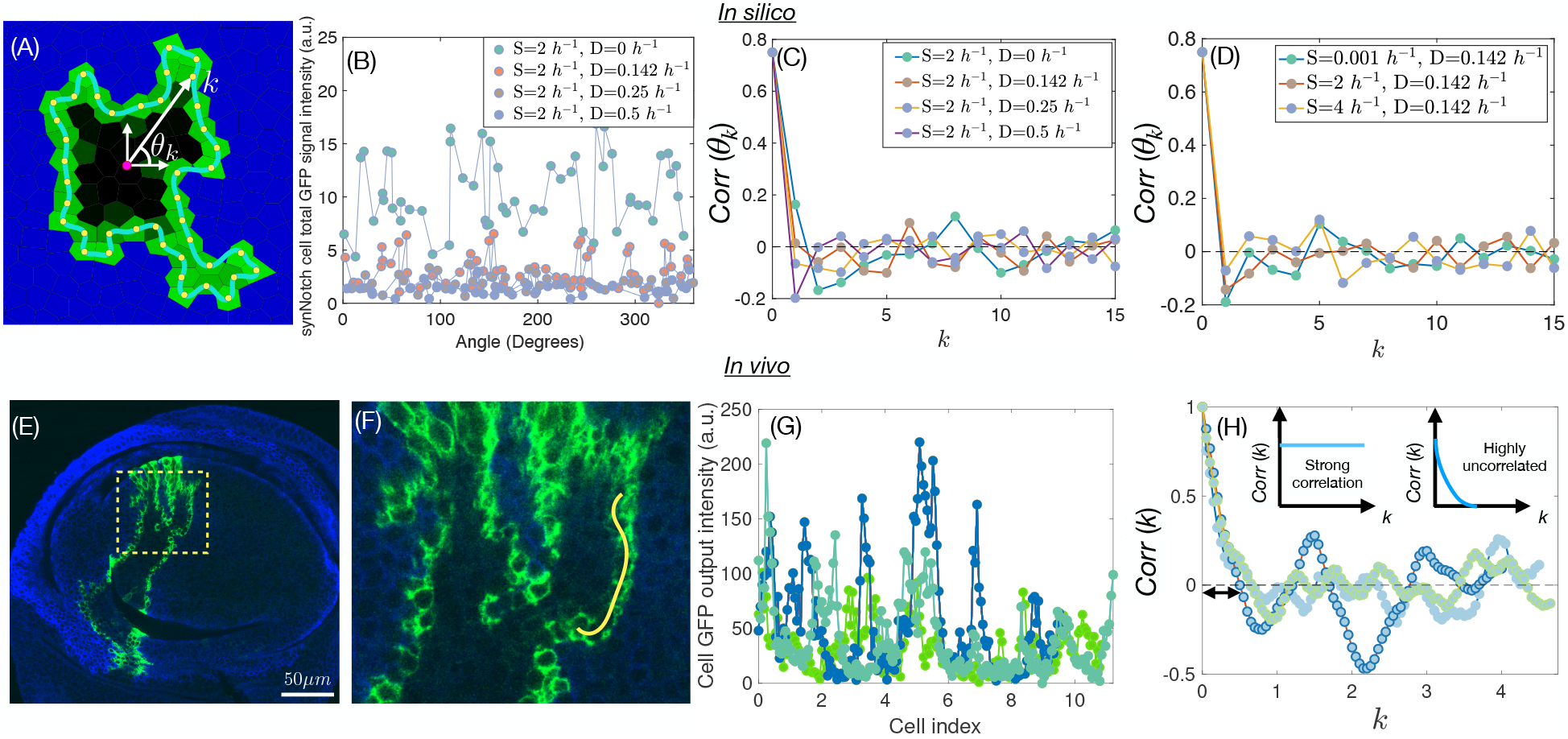
Output along the boundary of synNotch cell clone is highly variable and spatially uncorrelated. **(A)** Profile of output in border synNotch cells directly adjacent to ligand cells was quantified at t = 80h and geometrically fixed across multiple simulations using the angular position (k) of a synNotch cell relative to the synNotch clone center (see Materials and Methods). **(B)** Output intensity in the border synNotch cells for fixed synthesis rate (S) and various values of degradation (D).**(C**,**D)** Autocorrelation function of the output intensity in the border synNotch cells for (C) fixed *S* = 2*h*^−1^ and varying *D*, and for (D) fixed *D* = 0.142*h*^−1^ and varying *S*. Fast decay in the output intensity fluctuation correlation is characteristic of the high variability in the output among synNotch cells at the border. **(E**,**F)** Wing disc with clones of ligand (blue) and synNotch cells (uncolored), and GFP output (green) intensity measured along the yellow line across the border synNotch cells (F). **(G)** Output intensity relative to the typical cell size of 3μm along the border at 3 positions of the synNotch clone. (H) Autocorrelation analysis of fluctuations in the output intensity along the border rapidly decays to zero. This indicates large fluctuations in output intensity (inset plots show correlation as a function of cell index when there is strong correlation in output versus a highly uncorrelated scenario).

We quantified the output heterogeneity along boundary cells using the GFP intensity angular correlation function,

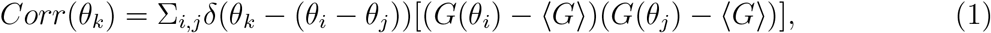

where *G*(*θ*_*i*_) is the GFP intensity in a cell whose angular position is at *θ*_*i*_ and *i, j* = 1…*N*_*b*_ are the indices for the total number of boundary cells (*N*_*b*_). *δ*(*z*) = 1 if *z* = 0, and 0 otherwise. ⟨*G*⟩ is the mean GFP intensity along the boundary. The decay in the correlation function quantifies the similarity in the GFP output between any two synNotch cells along the boundary, with quicker (slower) decay denoting low (strong) correlation. We found that the correlation function decays rapidly with changing angle or equivalently distance along the boundary irrespective of *D* (Fig. 4(C)) or output synthesis rate (*S*) (Fig. 4(D)). This implies that cell-to-cell variations in the GFP output at the synNotch clone border is independent of both *S* and *D*.

Next, to verify the above predictions we measured the GFP output intensity in synNotch cells *in vivo* along a section of the ligand-synNotch boundary (Fig. 4(E-F)). We found the GFP intensity profiles to be quantitatively similar to the simulations, with large fluctuations along the border of the synNotch clone, Fig. 4(G). Note that there is a difference in the GFP quantification between the simulation and *in vivo*: while we track GFP output per cell *in silico*, in the experiments we measure it at the cell border since the GFP is expressed at the cell surface. But, as we average over a line of finite thickness in the *in vivo* quantifications, we were able to make a direct comparison with the simulations. Our autocorrelation analysis confirmed that the GFP output correlation decays rapidly along the boundary, as predicted in the model (Fig. 4(H)). Therefore, we confirm that the spatially uncorrelated synNotch output patterns are a feature of synNotch output in growing tissues. An important consequence of cell-cell contact length dependent output synthesis is that differences in contact length between individual cells at the boundary can lead to highly variable juxtacrine signal output patterns.

### synNotch output decays exponentially and extends up to 3 cell diameters from the clonal ligand-synNotch interface *in vivo*

As juxtacrine ligands are membrane anchored their signaling is restricted to receptors on the surface of neighboring cells with which they make direct contact^45^. However, a feature we observe both *in silico* and *in vivo* is that the synNotch ouput is found several cell diameters from the ligand-synNotch interface. Our model points to the output degradation rate (*D*) as an important factor in determining the spatial gradient of the GFP output across the synNotch cell clone. In the model, we vary *D* from 0 − 0.5*h*^−1^ including the physiologically relevant value of *D* ∼ 0.142*h*^−1 46,47^ keeping the output synthesis rate (*S*) fixed at 2*h*^−1^. We observe generally that the border synNotch cells in contact with the ligand cells have high intensity of output (bright green), which gradually weaken (dark green to gray) towards the center of the clone (SI Fig. S2(A-B)) - a spatial pattern strongly dependent on *D*. For a low degradation rate, even synNotch cells not directly at the clone border have high output levels (see SI Fig. S2(A)). On the other hand, at high degradation rate cells have little to no output (gray) towards the clone interior and high output levels at the periphery (see SI Fig. S2(B-C)). Hence, the extent to which GFP output is lost in the synNotch clone interior is largely determined by *D*.

To quantify the GFP output gradient, we focused on the spatial region traversing the boundary and interior of the simulated synNotch clone in Fig. 5(A). The spatial output profile peaks at the clone boundaries and descends toward the clone interior (see Fig. 5(B)). With increasing *D*, the output decay is steeper (Fig. 5(C)). Moreover, the output intensity normalized to the synNotch cell at the interface (Fig. 5(C)) is well described by an exponential function of the form 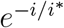, where *i* is the cell index and *i*\* is the characteristic number of cells from the border over which the output amplitude decays by 1*/e*. For *D* = 0, *i*\* ∼ 2, implying that a significant drop in output intensity occurs beyond 2 cells from the clonal interface. For *D* = 0.5*h*^−1^, *i*\* ∼ 0.5, indicating that the output intensity decays substantially within one cell layer from the boundary (see Fig. 5(D)).

**FIG. 5:**
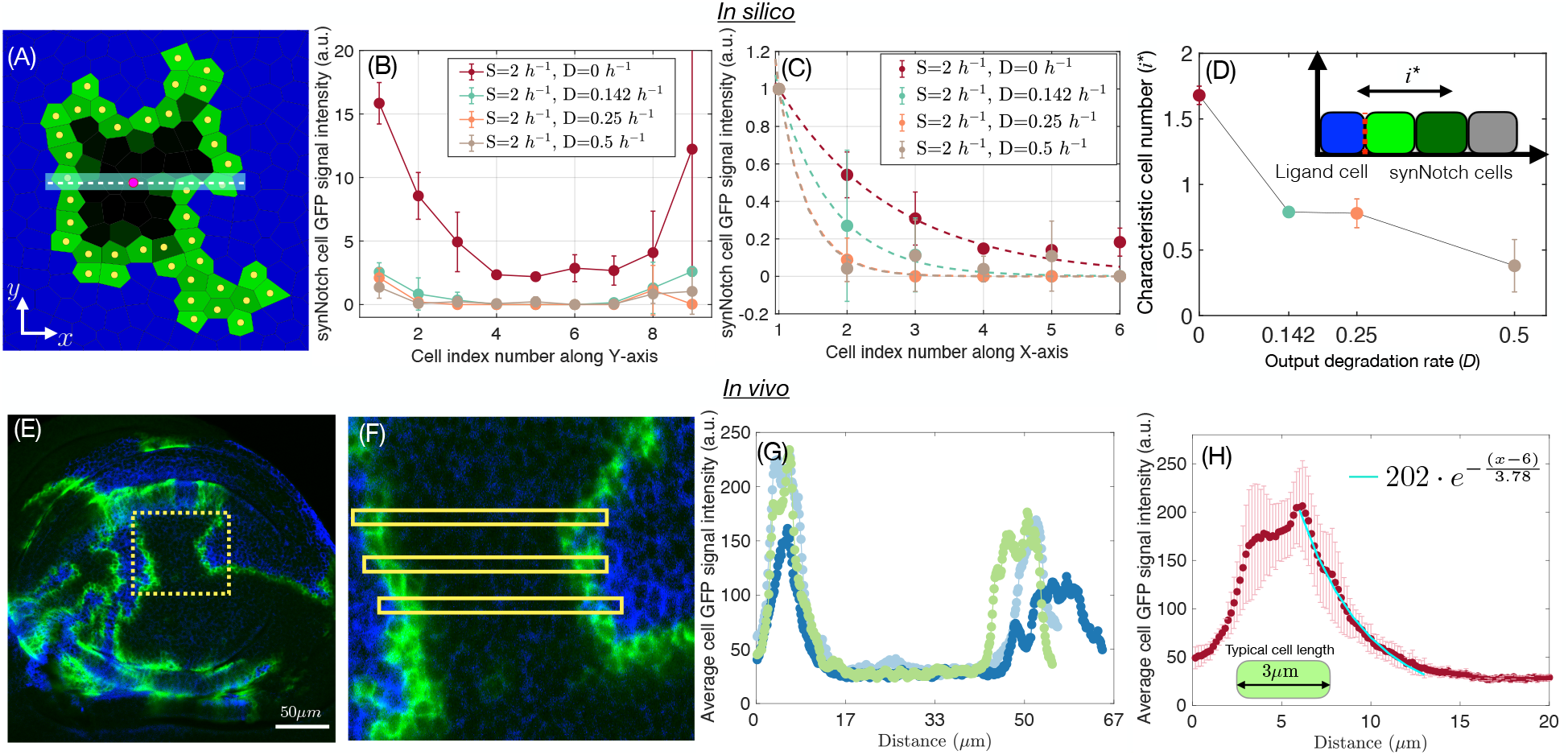
synNotch output decays exponentially away from the clonal interface depending on the output degradation rate. **(A)** The output across a simulated synNotch clone at t=80h is measured at the dashed line crossing the clone center which is geometrically fixed amongst multiple simulations (see Materials and Methods). **(B)** GFP output profile across synNotch clones (3 simulations) for fixed output synthesis rate (S) and varying output degradation rate (D, inset). **(C)** Output shown in (B) relative to the synNotch cell immediately adjacent to the clonal interface. This rescaled data is fit by an exponentially decaying function (dashed lines) of the form 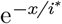. **(D)** The distance the output extends from the interface (i*) decreases with D. **(E**,**F)** Wing disc with clones of ligand (blue) and synNotch cells (uncolored), and GFP output (green) intensity measured within horizontal sections across the clone (yellow boxes). **(G)** Output profile at the 3 locations across the synNotch clone indicated in F. **(H)** The average of the three output profiles near the interface (magenta circles) fits an exponential function of the form 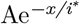 where i* = 3.78 μm (turquoise colored line). The average diameter of a synNotch cell is 3μm (inset). For analysis of multiple wing discs that show graded profiles see SI Fig. S3. Error bars=standard deviation.

Next, we compared our model predictions with synNotch activation *in vivo* using the GFP intensity in synNotch cells as a function of distance away from the ligand cells. Measurements across a single receptor clone showed a clear peak in fluorescence intensity at the boundary and lower intensity towards the clone center, Fig. 5(E-F). This is similar to our *in silico* predictions (compare Figs. 5(B) and (G)) and as a further quantitative test we found that the decay in the average GFP intensity is well described by an exponential function, Fig. 5(H). The fit revealed that the spatial gradient of GFP extends an average distance of ∼ 1.5 cell diameters and a maximum of 3 cell diameters.

To analyze this further, we selected clones with clear GFP expression in synNotch cells not adjacent to ligand cells and measured the spatial GFP expression profile (SI Fig. S3). This was done by fitting an exponentially decaying function (as in Fig. 5(H)) to the measured GFP intensity from the interface to the location where the GFP intensity profile plateaued at low values (Fig. S3(A4-E4)). We obtained the characteristic decay length of the GFP output into the clone from each *in vivo* experiment (SI Fig. S3(A4-E4)) and found that the output extended as far as 3 cell diameters away from the clonal interface (in 3 out of 5 experiments) and on average it was detected 2 cell diameters into the receptor clone (Fig. S3(F)).

In principle, we would expect the output from a juxtacrine signal to be strictly confined to cells that are directly in contact with the signal-sending ligand cells. Instead we found a spatially graded output that on average extends multiple cell diameters from the boundary into the receptor clone. Importantly and in contrast to the variation along the clonal interface that we show depends on contact length at all rates of output degradation (above), spread of the output into the clone interior is highly dependent on the rate of degradation of the output. Therefore our combined approach has identified two separate mechanisms acting to profoundly shape the overall pattern of a juxtacrine output at the tissue level.

## DISCUSSION

In this work, we tackled the complexity underlying signaling *in vivo* by taking a reductionist approach through engineering a synthetic signaling system based on the Notch receptor (synNotch) that we show operates orthogonally to native signaling systems in the *Drosophila* wing disc. We predicted the complex emergent patterns of output in clonal populations of ligand and synNotch wing disc cells by implementing a minimal computational model based on the hypothesis that synNotch output will depend on cell-cell contact length. We found that the synNotch output in the expanding tissue forms complex patterns and suggest that the major parameters that determine these patterns are: (i) contact length dependent syn-Notch activation, (ii) output synthesis and degradation rates and (iii) cell area growth and division.

### Ligand-synNotch cell contact area largely determines signal output

We found that the contact area of the interface between ligand and synNotch cells is an important determinant of the output pattern. We present evidence, consistent with previous results^40^, that this is due to the linear relationship between cell-cell contact area and the extent of receptor activation. We found that the output intensity is characterized by rapidly decaying spatial correlations and strong fluctuations along the ligand-receptor clonal interface. Our model predicted that this behavior arises due to heterogeneity in the interface contact lengths between synNotch and ligand cells, confirmed by the agreement with *in vivo* quantifications.

We identified two prominent sources of contact length heterogeneity: (i) a stochastic cell area increase as individual cells grow, and (ii) the random cell division orientations. As the cell division plane is assigned stochastically, a responding daughter cell may be displaced such that it either completely loses contact with the ligand cells or maintains contact, but most often with a smaller contact length. We conclude that these cell division outcomes must lead to marked variability in the extent to which synNotch is activated in the daughter cells and likely accounts for variability in output along the interface. Another consequence of cell division is that we expect the signaling interface to become increasingly irregular with growth. We therefore predict that extensive tissue growth will add points of geometric inflection to the interface, leading over time to greater variability in both contact area and the strength of the juxtacrine signal output.

In proliferating tissues where cell division axes are oriented randomly, as in the fly wing^48^, we assume that the above phenomenon is relevant, meaning that there will be emergent variations in a juxtacrine signal output. Presumably native developmental juxtacrine pathways have elements that compensate for this marked variation. This variation would also need to be factored in when engineering synthetic juxtacrine pathways, for instance to produce a persistent zone of cells that are required to respond uniformly as the tissue continuously proliferates. Feedback circuits might be one way to produce a precise region of persistent response following an initial signal activation. Positive feedback is common to many signaling pathways, including in native Notch activation, for example during the formation of the boundary between dorsal and ventral cell populations in the wing^49^. Future studies could produce models incorporating feedback circuits to rapidly determine which is most effective in dampening the variation in our system and to highlight components of native pathways that might counter variability due to cell division to form sharp borders between distinct cell populations. In cases where the synNotch response itself affects proliferation and morphology, a complex feedback would be expected between contact-area and receptor activation, necessitating the use of more sophisticated mathematical models to accurately predict the signal output at a particular time and location^50–54^.

In summary, while molecular scale variabilities such as temporal differences in the level of receptor activation may also impact synNotch output^55^, our results suggest that the dominant physical parameter that determines juxtacrine output *in vivo* is cell-cell contact area. As this is a function of the tension of the cell bonds between clonal ligand and synNotch cell populations^56^ and is regulated by the subcortical network of cell acto-myosin, our work also provides a foundation to study how mechanical forces generated by myosin and actin^57^, together with cell-cell adhesion molecules^58^ might regulate the spatial patterning of a juxtacrine signal output.

### The output of a juxtacrine signal can persist several cells away from the ligand/receptor interface and forms a gradient that decays exponentially

Naively, one would expect the juxtacrine signal output to be restricted to the responding cells strictly in contact with signal sending cells. Instead, we found that in the wing disc the output extends multiple cell diameters away from the ligand-synNotch interface in a graded output profile characterized by an exponentially decaying function. Our model replicates this profile in a manner dependent on output degradation rate, implying that at slower rates of degradation the output can persist and generate a long-range output. Specifically, when a responding synNotch cell divides, the output is equally shared between the daughter cells and at the same time a daughter cell may be displaced away from the clonal interface losing contact with the ligand cell. Multiple rounds of cell division, with the output decaying exponentially as each division halves the output, produces daughter cells retaining the output further from the interface, thereby establishing the graded profile that decays exponentially, as we observed *in vivo*.

Such graded profiles across a tissue are well defined for morphogens, such as Decapentaplegic (Dpp) and Wingless (Wg, a Wnt homolog), that diffuse from a population of source cells and elicit growth and transcriptional responses over 30 cell diameters away^59^ during normal developmental patterning^60,61^. Our observations suggest that juxtacrine signals might also have the capacity to generate similar graded profiles in development, providing the signal output is long lived and cell division sufficiently rapid. Interestingly, when Wingless is tethered or restricted to the surface of cells, transcriptional responses can still be detected in receiving cells several cell diameters away from the signal-sending cells^23^. This may arise due to a similar mechanism as we propose here - cell division-driven displacement of responding cells. Such activity might contribute to the surprising rescue of the development of animals where the only form of Wg is cell membrane tethered^62^.

Alternatively, it is possible that an output several cell diameters from the clonal interface may be induced by other means. For instance, basal cell extensions in the epithelium of the fly notum can support long-range Delta/Notch signaling over several cell diameters^63^ and there is evidence that cell extensions or cytonemes that extend basally across multiple cell lengths in the wing imaginal disc can act as signaling platforms for the morphogen Hedgehog^64,65^. However, it is unclear whether such extensions would convey a signal that decays from the apical cell interface with an exponential profile that is dependent on the rate of cell division. Further, our model does not include such long-range contact-based signaling, and restricts signaling to synNotch cells immediately adjacent to the ligand cells. Nevertheless, the model predicts graded exponentially decaying profiles that closely resemble those we observe *in vivo*. We cannot draw any conclusions about the ability of basal extensions to produce long-range juxtacrine outputs based on this study, but our model suggests that at least in principle perdurance of the signal response alone is sufficient to produce a long-range output. In the future, it will be interesting to more directly examine whether basal extensions have a role in the spatial profile we observe *in vivo*. The model will be an asset for this work since discrepancies between model and *in vivo* could indicate the possible influence of additional factors, like cytonemes, responsible for long-range output.

### Tools for the development of new synNotch and synthetic juxtacrine circuits

Our model is also the foundation for future tools to accelerate the design of synthetic cell-cell signaling circuits that will be of value in therapeutics, such as regenerative medicine and tissue engineering, where transcriptional responses need to be deployed at precise times and locations within a proliferating tissue. One immediate application of the model is to improve synthetic receptors by measuring the activity of new variants of synNotch. Comparison of a variety of synNotch receptors capable of producing a response from identical GFP transgenes, effectively fixing the output degradation rate, would allow the exponential decay of the output to be used to quantify the synthesis rate, which is proportionally related to the activation of the receptor. Further iterations of our approach, as discussed above, will also be of value in the design of more elaborate circuits to be deployed in complex tissue environments.

Overall, our work is an example of a combined strategy to expediate the design-buildtest-learn cycle of synthetic biology, using the model for the design phase and the fly wing to build and test synthetic tools, leveraging its proven capacity to decipher native signaling pathways.

## MATERIALS AND METHODS

### 1. *Drosophila* Method Details

#### *Drosophila* Transgenes

Fly stocks were maintained at 25^*o*^C on standard corn food, and all experiments were performed at 25^*o*^C on standard corn food. All ligand and receptor coding sequences, with the exception of FSH*α*, were inserted into a modified form of *pUAST-attB* (www.flyc31.org) that contains a single Flp Recombinase Target (FRT, *>*) positioned between the UAS promoter and the coding sequence, and the resulting *UAS > FSH* −*Delta* and *UAS > FSHR* − *synNotch* transgenes were inserted at a single genomic docking site, *attP* − 86*Fb* located on the right arm of the third chromosome and were oriented so that the promoter is centromere proximal to the coding sequence^66^. A single *UAS*.*FSHα*, transgene inserted by conventional P-element mediated transformation onto the X chromosome was used in all experiments. The FSH chimeric form of Delta is described in detail in Ref.^34^. Briefly, the native extracellular domain of Delta was replaced in its entirety by the *β* subunit of human Follicle Stimulation Hormone (FSH*β*)^67^ and an HRP tag was inserted immediately downstream of the C-terminal YCSFGEMKE sequence of FSH*β* and immediately upstream of the Dl transmembrane domain. In all experiments, FSH*α* was co-expressed with the FSH*β*-Dl protein to reconstitute FSH, which is an FSH*α*/FSH*β* heterodimer (for simplicity, we refer to the ligand as the FSH-Delta chimera and do not include the HRP tag in its designations). The FSHR-synNotch form of Notch is based on the FSHR-A2-Notch receptor described in detail^66^. Briefly the amino-terminal Epidermal Growth Factor (EGF) Repeat containing portion of the native extracellular domain of N was replaced by the ectodomain of FSHR. The extracellular domain was tagged by the insertion of mCherry just upstream of the juxtamembrane A2 domain. The NRR of Notch was replaced with the force sensitive region of von Willibrand Factor A2 domain. Previous work has shown that this domain acts as a force sensor in the same way as the NRR^66^. The additional change to FSHR-A2-Notch to form FSHR-synNotch replaced the cytosolic domain of FSHR-A2-Notch with the LexA-VP16 trascription factor^68^. The join between Notch and Lex A is as follows (the Notch sequence is in uppercase): VITGIILVIIALAFFGMVLSTQRKRsgppkaltarqqevfdlir. Complete DNA sequences for both the ligand and receptor are available on request.

#### Analysis of signaling between FSH-Delta and FSHR-synNotch expressing cells

Signaling between FSH-Delta and FSHR-Notch cells was established using Mosaic Analysis by Promoter Swap (MAPS; see^34^ for detailed description). In brief, mitotic recombination across the FRTs in cells transheterozygous for UAS>transgenes is induced in the presence of a *nub*.*Gal4* driver that acts in the developing wing. Upon the appropriate segregation of the recombined chromatid arms at the 4 strand stage, populations of FSH-Delta and FSHR-synNotch expressing cells were generated; the FSH-Delta expressing cells are marked by staining for the HRP epitope tag (blue in all of the Figures), and the dedicated FSHR-synNotch expressing cells are marked “black” by the absence of HRP expression. Receptor activation occurs at the interface between the cells that express FSH-Delta and those that express solely the receptor^66^. Here this was monitored by assaying for a GFP output, since FSHR-synNotch activation results in the translocation of the LexA-VP16 transcription factor to the nucleus and expression of a transgene composed of the LexO promoter followed by a fusion of the surface protein CD2 followed by GFP (LexAop rCD2::GFP, Bloomington Stock Center). To induce mosaics by promoter swap, first or second instar larvae of the appropriate genotype (*hsp*70.*flp UAS*.*FSHα/hsp*70.*flp* or *Y* ; *nub*.*Gal*4*/LexAop rCD*2 :: *GFP* ; *UAS > FSH* − *Delta/UAS > FSHR* − *synNotch*) were heat shocked at 33^*o*^C for 20 minutes and wing discs from mature third instar larvae were dissected, fixed, stained for HRP expression and mounted for confocal microscopy [commercially available Rabbit anti-HRP (Abcam ab34885, 1/1000) antisera were used in combination with Alexa 633 conjugated labelled secondary antisera following standard protocols, e.g., as in^34^]. The native fluorescence of the GFP was also visualized in these preparations.

### 2. Vertex based modeling framework for synNotch induced gene expression

We use the vertex model to represent the *Drosophila* wing disc epithelium as a two dimensional cell layer. Using a cell-based computational framework, Virtual Leaf^69,70^, we simulate coupled vertex and chemical dynamics we implemented. The two-dimensional description is a good approximation for the wing disc epithelium if changes in cell height are assumed to be negligible, Fig. 1(D). Cells in this epithelial monolayer are represented as a network of interconnected polygons (edges as straight lines connecting vertices). The configuration of each individual cell, and thus of the whole network, is characterized by the positions of the vertices, and the connections between them. Assuming cell shape relaxations occur at timescales much faster than the cell division, the stable network configuration is determined by the minima of the energy function,

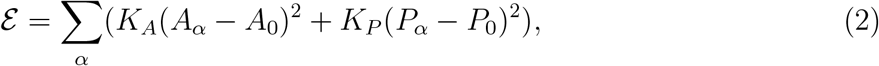

where, *A*_*α*_ and *P*_*α*_ are the area and perimeter of cell *α*, respectively. *A*_0_ and *P*_0_ are the preferred area and perimeter of cell *α*. The summation is over all cells *α* = 1 ⋯*N*(*t*), where *N*(*t*) is the total number of cells at time *t*. The first term describes the cell area elasticity with an elastic coefficient *K*_*A*_ which results from cell volume incompressibility and the cell monolayer’s resistance to height fluctuations. The second term describes the stiffness of the cell perimeter *P*_*α*_ with an elastic constant, *K*_*P*_, and it results from two processes: contractility of the actomyosin subcellular cortex and effective cell membrane tension due to cell-cell adhesion and cortical line tension. The energy function described by Eq. (2) is widely used to study tissue mechanics during morphogenesis^71,72^.

A detailed description of the VirtualLeaf vertex modeling framework, which we have adapted here, is given in Ref.^69^. The shape of a cell, and thus the whole tissue, is determined by the force balanced state, i.e., the state in which the forces, described in the energy function E given by Eq.(2), acting on each node have sufficiently balanced out. The simulation of this vertex model employs Monte Carlo based ‘Metropolis dynamics’ method to alter the positions of the vertices until the energy function, given by Eq.(2), is minimized such that all interfacial tensions are balanced by cellular pressures^69^. We assume the mechanical properties of all the cells of both the cell types in the tissue, i.e., ligand and receptor-expressing synNotch cells, to be homogeneous and constant. The values of the model parameters are listed in the Table (I).

**TABLE I:**
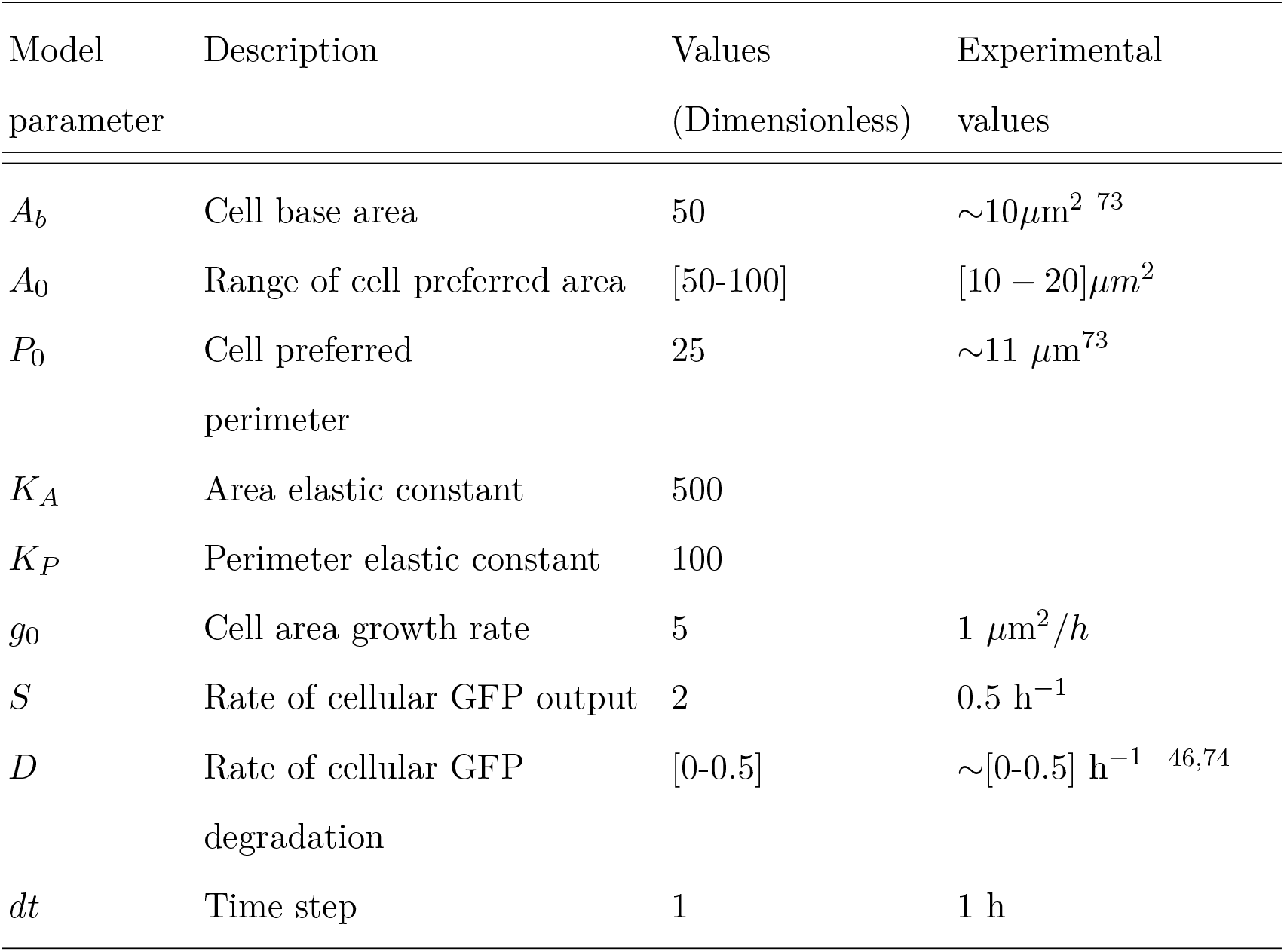
Model parameters, their meaning, dimensionless values and equivalent magnitude in experimental units.

| Model<br>parameter | Description | Values<br>(Dimensionless) | Experimental<br>values |
| --- | --- | --- | --- |
| $A_b$ | Cell base area | 50 | $\sim 10 \mu\text{m}^2$ <sup>73</sup> |
| $A_0$ | Range of cell preferred area | [50-100] | $[10 - 20] \mu\text{m}^2$ |
| $P_0$ | Cell preferred<br>perimeter | 25 | $\sim 11 \mu\text{m}^{73}$ |
| $K_A$ | Area elastic constant | 500 | |
| $K_P$ | Perimeter elastic constant | 100 | |
| $g_0$ | Cell area growth rate | 5 | $1 \mu\text{m}^2/h$ |
| $S$ | Rate of cellular GFP output | 2 | $0.5 \text{ h}^{-1}$ |
| $D$ | Rate of cellular GFP<br>degradation | [0-0.5] | $\sim [0-0.5] \text{ h}^{-1}$ <sup>46,74</sup> |
| $dt$ | Time step | 1 | 1 h |

### 3. Dynamics of GFP signal pattern in synNotch cells

To characterize the spatiotemporal pattern of synNotch induced GFP output in the *Drosophila* wing disc, in addition to the dynamics of cell shape and cell division, we incorporate a dynamic equation for the change in the number of GFP molecules in a synNotch cell. The cell GFP signal intensity at any time *t* in our model is proportional to the total number of GFP molecules in the cell at that time. Our modeling framework thus gives rise to a coupling between three timescales at the scale of a single cell, namely, the intracellular GFP synthesis rate, the rate of intracellular GFP degradation and the rate of cell division.

In our model, the wing disc epithelium consists of two types of cells, namely, synthetic ligand expressing cells and synthetic Notch receptor expressing cells (which we refer to as synNotch cells). The ligand expressing cells are assigned a label or cell type of 0 and the synNotch receptor cells are assigned a label or cell type of 1. This binary-type quantity *T* is a function of cell *α*, where *α* is the cell index, *i* ∈ 1 … *N*. *T* takes the value *T*(*α*) = 0 if the cell *α* is a ligand expressing cell and *T*(*α*) = 1 for a synNotch receptor cell, Fig. 1(A). The total number of cells at any given time point is *N*(*t*) = *N*_(*T* =0)_ + *N*_(*T* =1)_, where *N*_(*T* =0(1))_ is the total number of cells of type *T* = 0(1). We assume that the whole wing disc is fully characterized as comprising of type 0 and type 1 cells. The cell identity, *T*(*α*) = 0 or 1, assigned to a cell is conserved across cell division events. Therefore, when a cell *α* of type *T*(*α*) = 0, i.e., a ligand expressing cell, divides, it produces two daughter cells, both of which are of cell type 0. Similarly, when a cell of cell type *T*(*α*) = 1, i.e., a synNotch cell, divides, it produces two daughter cells, both of which are cells of cell type 1. We propose that the dynamics of GFP output (*G*_*α*_) as a function of *t* is given by,

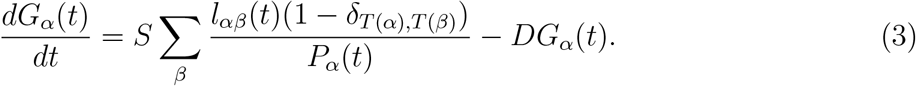

The first term on the right hand side of Eq.(3) describes GFP synthesis at the rate *S*(*h*^−1^) in synNotch cells. The summation is over all cells *β* neighboring a cell *α*. GFP is synthesized only in those synNotch receptor cells that are adjacent to the synthetic ligand expressing cells, i.e., synNotch receptor cells which share an edge or a cell bond with the ligand expressing cells. This condition is ensured in the model by the kronecker delta function *δ*_*T*(*α*),*T*(*β*)_, which assumes the value 0 if the Notch cell *α* and its adjacent cell *β* are of different types, i.e., (*T*(*α*) ≠ *T*(*β*)), and, the value 1 if they are of the same type, i.e., (*T*(*α*) = *T*(*β*)). With this condition, GFP synthesis is restricted to only those cells that share a cell-cell junction with synthetic ligand expressing cells. We propose that GFP output in a synNotch cell is proportional to the length of the cell bond, *l*_*αβ*_ (*μ*m) shared with its adjacent synthetic ligand expressing cell, and normalized to the overall perimeter *P*_*α*_ (*μ*m) of the synNotch cell *α* at time *t*. The second term on the right hand of Eq.(3) accounts for the degradation of GFP in the synNotch cell *α* at a constant rate of *D* (h^−1^). Now, we briefly describe the non-dimensionalization of Eq.(3). By multiplying both sides of the Eq.(3) by *t*\* = 1*h* we can write the dimensionless form of Eq.(3) as,

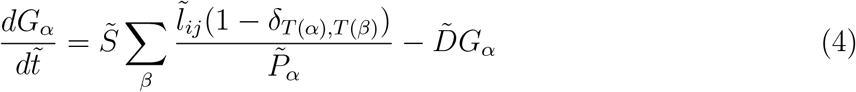

where, 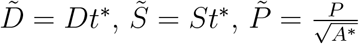 and 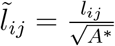.

### 4. Analytical derivation showing dependence of the total synNotch GFP expression on the shape of the synNotch cell colony

The time dependent dynamics of the total GFP expression in the whole synNotch cell colony is described by the following equation,

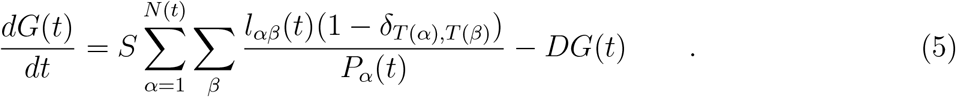

If we assume each individual cell within the synNotch cell colony has roughly the same perimeter which is 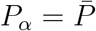 then,

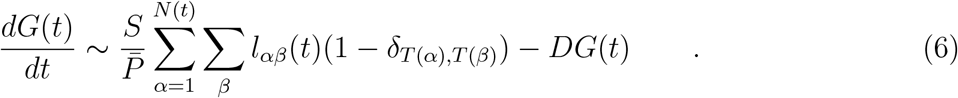

Let us introduce an average cell shape factor 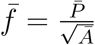, where 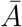 is the average single cell area and is defined as 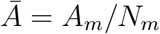, with *A*_*m*_ being the total area of the whole synNotch cell colony, and *N*_*m*_ being the total number of synNotch cells in that colony in the time interval *t* ∈ [*τ*_*m*_, *τ*_*m*+1_] between two succesive cell division events.

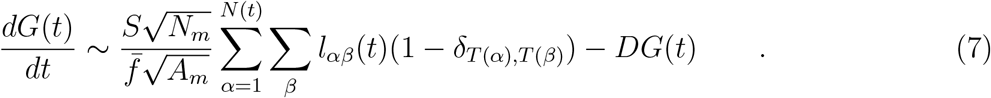

The double summation in Eq. (7) actually results in the perimeter of the whole synNotch cell colony in the time interval *t* ∈ [*τ*_*m*_, *τ*_*m*+1_], which we will denote by *P*_*m*_.

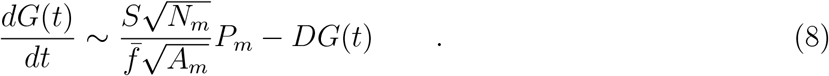

The factor 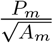 in Eq. (8)is the shape factor of the whole synNotch cell colony at time *t* which we will denote *F*_*m*_.

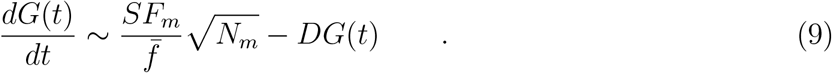

The above equation can also be written in terms of circularity of the whole synNotch clone. The circularity parameter is related to the shape factor in the following manner, 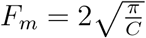,

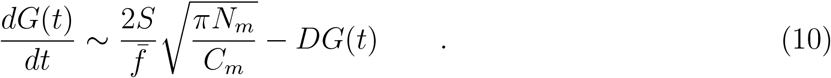

Therefore, we predict that when the circularity of the synNotch clone is small (at lower values of *C*_*m*_), we expect the source term contribution to be effectively larger. This would lead to an enhanced synNotch output.

### 5. Rules governing cell growth and division

Each cell in the *in silico* tissue is fixed to the same base area, *A*_*b*_. Our initial condition is such that all the cells are set to *A*_*α*_ = *A*_*b*_ = 50, where *α* is the cell index. At *t* = 0, the preferred area *A*_0_ of all the cells is equal to the base area. The dimensionless value of 50 for the cell area in the model corresponds to an experimental value of 10*μ*m^2^. To implement cell area growth over time, as described in^69^, the cell preferred area is increased in the next time step *t* + *δt* as,

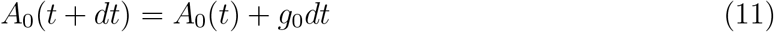

where, *g*_0_ is the value by which the cells preferred area is incremented in the next time step. The cell area at each time step is relaxed quasi-statically to the preferred area, which results in an irreversible increase in the cell area over time. Over time, when the area of a cell *A*_*α*_ is double the base area *A*_*b*_ then the cell divides over its short axis, Fig. 1(D-iv). Right after division, the preferred area of the divided cell is reset to *A*_0_ = *A*_*b*_ = 50. As a result the area of each daughter cell, the instant after cell division, is relaxed to *A*_0_ = *A*_*b*_. Since the cell divides when its area *A*_*α*_ reaches a value which is twice the base area of the cell *A*_*b*_, which occurs in the time duration *τ*,

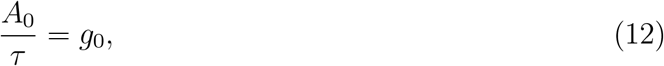

in our simulations we chose a timescale such that 1 relaxation step that minimizes the energy function *E*, given by Eq. (2), for the preferred *A*_0_(*t*) at the time *t*, corresponds to 1 h. We set *A*(*t*) = *A*_0_, where *A*_0_ = 10*μm*^2^ is the typical base area of a cell in the wing disc epithelium. The typical time *τ* over which a wing disc epithelium cell divides is equal to 10 h. Hence, *g*_0_ = 1*μm*^2^*/h* is the typical cell area growth rate. By dividing and multiplying both sides of Eq.(12) by a unit area *A*\* = 0.2*μm*^2^ and a unit time *t*\* = 1*h*, respectively, we obtain,

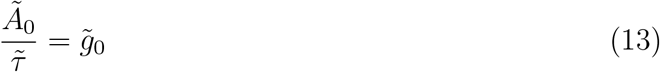

We chose the cell base area as 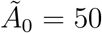, which corresponds to the experimental value of 10*μm*^2^, and cell division time 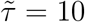, which is equivalent to 10h, implying that 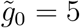.

### 6. Bio-image analysis

For measuring the cell perimeter and the corresponding synNotch output (GFP) from experimental images, we used the free and open-source bioimage analysis software CellProfiler^43^ and CellProfiler Analyst^44^. The provisions within CellProfiler for automated cell segmentation was used to identify the cell membrane borders of individual cells in images that we analyzed. The difference in the GFP intensity between the bright cell membrane of a responding cell and the corresponding dark cytoplasm of the cell interior was used to segment out the cell membrane of each of the identified individual synNotch cells. Intensity of GFP response along the cell membrane was measured for each cell together with the cell shape parameter quantifications. These measurements were then filtered using CellProfiler Analyst, where a machine learning algorithm selected the misidentified cells. The CellProfiler Analyst was imperfect in its ability to completely filter out all misidentifications, so further manual filtering was required to obtain the results shown.

### Parameter values

## ACKNOWLEDGMENTS

A.M.K acknowledges support from the College of Science and Mathematics at Augusta University (AU). P.L. acknowledges support from NICHD (R21HD107414), the Roy and Diana Vagelos Precision Medicine Award and the Research Scholarship & Creativity Activity Program at AU. P.L. thanks Gary Struhl for assistance with transgene design and Olivia Mosley for technical support. A.M.K and P.L. acknowledges support from the AU Center for Undergraduate Research and Scholarship (CURS) Summer Scholars Program. We thank Jessica Hoffmann, Sumit Sinha, Xin Li and Eric Vitriol for their valuable comments on the manuscript.

## Notes

### Competing Interest Statement

The authors have declared no competing interest.

### Summary of Updates

Results section shortened and updated to better focus on the main results; New revised Figure 3; Supplemental files updated.

