## Supplementary Information for "Contact area and tissue growth dynamics shape synthetic juxtacrine signaling patterns"

(Dated: 7 May 2024)

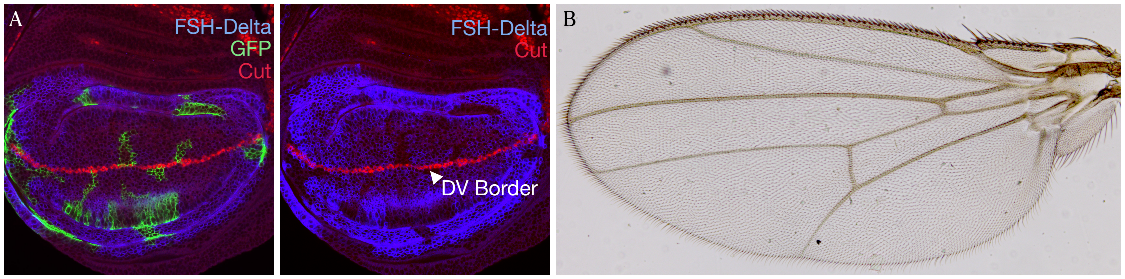


FIG. S1: FSHR-synNotch does not crosstalk with or inhibit endogenous Notch. A) Representative third instar wing imaginal disc with FSHR-Notch clones meeting FSH-Delta clones (marked in blue by anti-HRP antibody staining). At the clonal interface, the FSHR-synNotch target gene GFP (green) is expressed (as in Fig 1B). During normal development, the Notch target gene cut (shown in red) is expressed at the border between dorsal and ventral cells (DV border). The overexpression of FSH-Delta and FSHR-synNotch does not produce ectopic Cut nor perturb endogenous Cut expression at the DV border, indicating there is no interaction between the synNotch and endogenous Notch signaling events. B) Representative adult wing from a fly expressing FSHR-synNotch and FSH-Delta as in A. The wing appears wildtype and has no obvious developmental phenotype.

### I. FSHR-SYNNOTCH DOES NOT CROSSTALK WITH OR INHIBIT ENDOGENOUS NOTCH

It is possible that versions of synNotch could alter native Notch activation or that synNotch ligands have the capacity to activate other Notch receptors. If so, this would lead to ectopic growth and complicate the prediction of the time and location of a synNotch response, as well as limiting the use of multiple synNotch receptors within the same system. Previous work has shown that FSH-Delta does not lead to the expression of Cut in neighboring wing cells that possess endogenous Notch and do not express the cognate FSHR-Notch receptor[1]. Similarly, FSHR-Notch shows no activation when presented with native Notch ligands[1]. Fig. S1 shows that there was no sign of native Notch activation (as indicated by Cut expression) at the FSH-Delta/FSHR-synNotch clone borders, despite the activation of FSHR-synNotch. This indicates that FSHR-synNotch functions orthogonally to native Notch/Delta. This is consistent with other synNotch studies that have demonstrated that multiple synNotch pairs can operate orthogonally within the same cell[2,3]. For a more stringent test, the phenotype of adult wings expressing synNotch proteins was examined. Even slight ectopic Notch activation can induce ectopic wing margin in the adult, whilst Notch inhibition results in wing notching. As shown on the right, a wing expressing both FSH-Delta and FSHR-synNotch developed normally to produce a wildtype adult wing. Therefore, FSH-Delta/FSHR-synNotch operate orthogonally to native Notch and its ligands.

### II. SIMULATION MOVIES

Movies of synNotch signaling in growing *in silico* epithelial tissue. The total duration of the tissue growth is 80 hours (h).

**Movie 1:** In silico synNotch signaling in a growing epithelial tissue for output synthesis rate ($S = 2/h$ and output decay rate $D = 0/h$ (Link)

**Movie 2:** In silico synNotch signaling in a growing epithelial tissue for output synthesis rate ($S = 2/h$) and an intermediate output decay rate $D = 0.25/h$ (Link)

**Movie 3:** In silico synNotch signaling in a growing epithelial tissue for output synthesis rate ($S = 2/h$) and higher output decay rate $D = 0.5/h$ (Link)

### III. SPATIAL OUTPUT PATTERN IN SYNNOTCH CLONES DEPENDS ON THE DEGRADATION RATE $D$

To further understand the role of the degradation rate in determining the synNotch output, we examined the spatial distribution of GFP output within the synNotch clone. We observe generally that the border synNotch cells adjacent to ligand cells have high intensity of GFP output (bright green), which gradually weaken (dark green to gray) towards the center of the clone (Fig. S2(A-C)). The lower intensity of the output at the clone center could be due to the absence of GFP synthesis in synNotch cells that lose contact with the ligand cells, yet continue to degrade the output over time. Accordingly, we expect the GFP intensity within the synNotch clones to be strongly dependent on the degradation parameter $D$. For small $D$, representing a slower output decay rate, output remains throughout the whole clone with little variation (Fig. S2(A)). At the intermediate value of $D = 0.25h^{-1}$ in Fig. S2(B), a “donut”-like pattern emerges with little to no GFP (dark green to gray) in cells at the clone interior while cells at the periphery have higher output levels. At even higher values of the output decay rate ($D$), the “donut”-like pattern is preserved but output in the peripheral cells is also lower, leading to more marked heterogeneity in output levels amongst cells (see Fig. S2(C)). Taken together, our results suggest that (i) the total GFP output in the synNotch clone is determined not only by the total number of synNotch cells but also by the clone shape, (ii) GFP output pattern within the clone is determined by

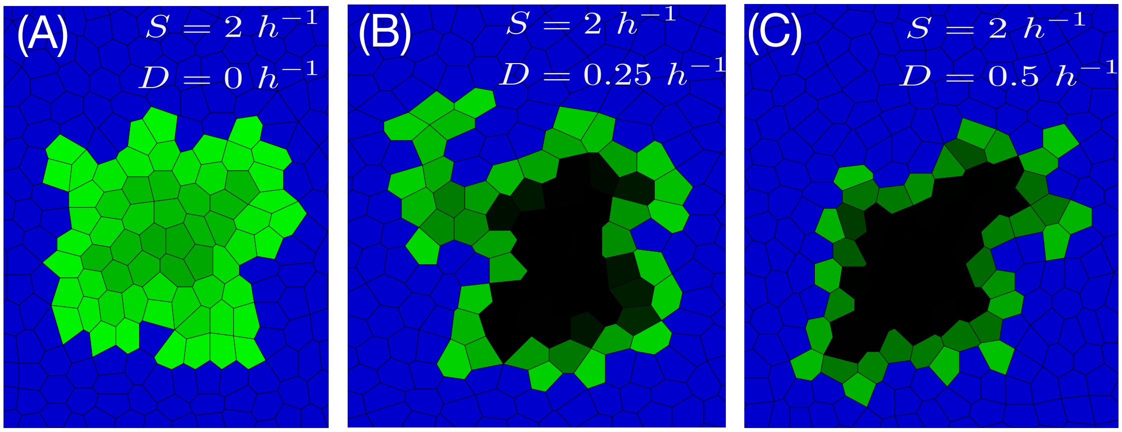


FIG. S2: **The degree of synNotch output spread into the synNotch clone is dependent on the degradation parameter (D). (A-C)** Pattern of output intensity in synNotch cell populations after $t = 80h$ for fixed synthesis rate (S) and varying signal output decay rates (D).

the intracellular GFP output degradation rate. Of note, some synNotch cells *in silico* show significant output despite not directly contacting ligand cells, which we investigate further in the main text and below.

### IV. SYNNOTCH ACTIVATION IN GROWING TISSUES FORM A SPATIAL OUTPUT GRADIENT

We found evidence for a spatial gradient of output extending from the signal-sending cell interface, which may be driven by the interplay between clone shape and $D$. We analyzed *in vivo* examples of signaling interfaces where we detect GFP output beyond the ligand/synNotch interface, and quantified the output profiles as a function of distance from the interface. To appropriately mark the clonal interface, anti-HRP antibody staining (Fig. S3(A1-E1)) against the HRP tag of the FSH-Delta ligand is used. This effectively marks the ligand cell populations and hence the interface between receptor and ligand cells (see yellow line in Fig. S3(A1)). Similarly, the output intensity is obtained as shown in Fig. S3(A2-E2). The expression profiles of both HRP staining (red line) and GFP (green line) along a horizontal line in the region of interest traversing both ligand cells and synNotch cells are shown in Fig. S3(A3-E3).

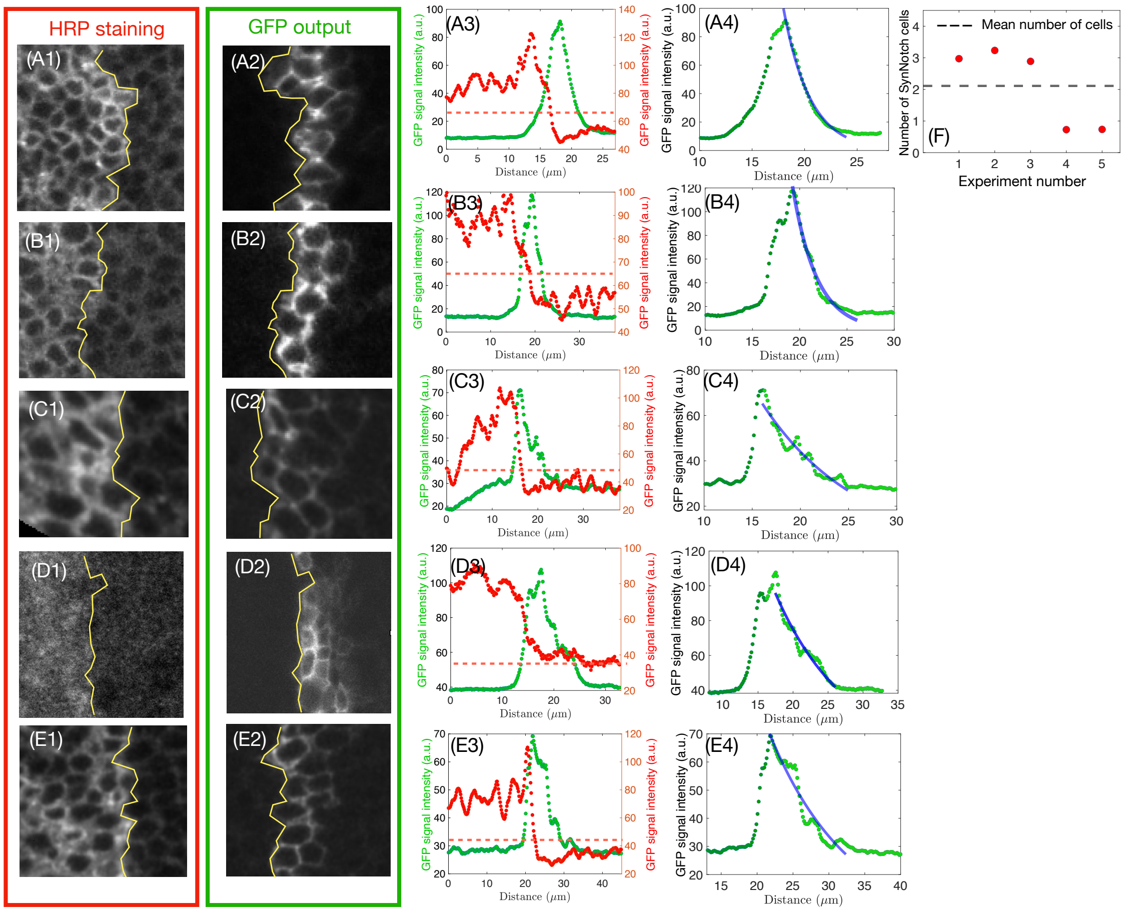


FIG. S3: **GFP output can extend beyond the ligand synNotch interface as far as 5 cell diameters. (A1-E1, A2-E2)** Five distinct ligand/synNotch interfaces where HRP staining (A1-E1) marks ligand-expressing cells to clearly identify the location of the clonal border (yellow line, A1-E1), allowing quantification of the spread of GFP output (A2-E2) into the synNotch cells. **(A3-E3)** GFP fluorescence intensity was measured along a straight horizontal line passing through the interface between synNotch and ligand cells (red dots=HRP staining intensity, green dots=GFP output intensity). For quantification we designated the precise interface location between the ligand and synNotch receptor cells as the distance where the fluorescence intensity of the HRP dropped to 45% of the maximum (shown by dashed red lines). . The intensity of the GFP signal output at this location is labeled $G_{max}$. **(A4-E4)** Across multiple experiments, the output decays exponentially from the signal interface. The exponential function $G_{max}e^{-x/x^*}$ (solid blue curve) fit the intensity profile of the output from the location of $G_{max}$ to the point where the output plateaus. **(F)** Quantification from each experiment of the number of cells from the clonal interface within which the output fluorescence is detected. We extract the number of cells from the interface and into the clone across which the GFP intensity drops by the factor $1/e$. We use the experimental scale wherein approximately $3\mu m$ corresponds to one cell diameter. The dashed line corresponds to the mean indicating spatial output propagation over 2 cell diameters.
